# Neural mechanisms underlying the effects of cognitive fatigue on physical effort-based choice

**DOI:** 10.1101/2024.12.06.627274

**Authors:** Michael Dryzer, Vikram S Chib

## Abstract

Fatigue is a state of exhaustion that influences our willingness to engage in effortful tasks. While both physical and cognitive exertion can cause fatigue, there is a limited understanding of how fatigue in one exertion domain (e.g., cognitive) affects decisions to exert in another (e.g., physical). We use functional magnetic resonance imaging (fMRI) to measure brain activity while human participants make decisions to exert prospective physical effort before and after engaging in a cognitively fatiguing working memory task. Using computational modeling of choice behavior, we show that fatiguing cognitive exertion increases participants’ subjective costs of physical effort compared to a baseline rested state. We describe how signals related to fatiguing cognitive exertion in the dorsolateral prefrontal cortex influence physical effort value computations instantiated by the insula, thereby increasing an individual’s subjective valuation of prospective physical effort while cognitively fatigued. Our results support the idea of a general fatigue signal that integrates exertion-specific information to guide effort-based choice.

## Introduction

Fatigue can be induced by both physically and cognitively effortful tasks, and it is often perceived that fatigue in one domain of exertion can influence feelings in another. For example, after a long day of cognitively draining grant writing, we might decide not to participate in a physically fatiguing after-work pickup soccer game. Recent studies have shown that fatigue will increase self-reported perceptions of effort (Greenhouse-Tucknott *et al*., 2020; Pageaux, 2014) and decrease willingness to exert (Chong *et al*., 2017; Hogan *et al*., 2020; Müller *et al*., 2021). Functional neuroimaging studies have implicated a network of brain regions, including the anterior cingulate cortex (ACC), insular cortex, and ventromedial prefrontal cortex (vmPFC), in both cognitive and physical effort-based decision-making (Hogan *et al*., 2019; Pessiglione *et al*., 2018; Shenhav *et al*., 2017; Westbrook *et al*., 2019) and shown that these regions are sensitive to fatigue state (Hogan *et al*., 2020; Müller *et al*., 2021; Wylie *et al*., 2020). However, there is a limited understanding of how fatigue in one exertion domain (e.g., cognitive) influences decisions to exert other types of effort (e.g., physical), and how the brain integrates information about different effort and fatigue modalities when making decisions to exert.

Previous behavioral experiments have shown that sustained cognitive and physical exertion increases perceptions of fatigue and is associated with decreased behavioral performance (Marcora *et al*., 2009; Pageaux, 2014; Pageaux and Lepers, 2016). Crosstalk between cognitive fatigue and physical performance has been observed, with mental fatigue impairing physical endurance and motor skill performance, as well as perceptions of effort and feelings of general fatigue (Eddy *et al*., 2015; Marcora *et al*., 2009; Moore *et al*., 2012; Pageaux, 2014; Pageaux and Lepers, 2016). However, these works found that cognitive fatigue did not impair maximal motor exertion, suggesting that fatigue may influence the affective processing of effort independently from actual exertion capacity. While several studies have examined the behavioral influence of cognitive fatigue on physical exertion, there is a limited understanding of the neurobiological mechanisms through which cognitive fatigue impacts physical decision-making and willingness to exert.

Studies of the neural basis of cognitive and physical effort-based decision-making suggest a domain-general encoding of prospective effort value by brain regions including the vmPFC, anterior insula, and ACC (Aridan *et al*., 2019; Chong *et al*., 2017; Hogan *et al*., 2019; Hogan *et al*., 2020; Lopez-Gamundi *et al*., 2021; Massar *et al*., 2018; Müller and Apps, 2019; Pessiglione *et al*., 2018; Westbrook and Braver, 2015; Westbrook *et al*., 2019). Beyond this common effort network, neuroimaging analyses have also implicated effort-specific brain regions related to exertion (e.g., physical exertion: premotor cortex, motor cortex, sensorimotor cortex (Hogan *et al*., 2019; Hogan *et al*., 2020; Müller and Apps, 2019); working memory cognitive exertion: dorsolateral prefrontal cortex (Barbey *et al*., 2013; Westbrook *et al*., 2019)).

Recent theoretical and experimental studies have begun considering how fatigue impacts effort-based decision-making (Hogan *et al*., 2020; Müller *et al*., 2021; Renfree *et al*., 2014). These works have shown that fatigue inflates the subjective value of effort and makes individuals less willing to accept options associated with higher effort. Neuroimaging and behavioral modeling of effort-based choice revealed that frontal cortex and insular cortex represent physical fatigue states while individuals make effort-based decisions. It has been suggested that information related to bodily state could modulate decisions to engage in physical activity (Hogan *et al*., 2020; Stephan *et al*., 2016). This information may be integrated by brain regions responsible for value-based decision-making during choices to exert. These previous studies focused on how physical fatigue influenced physical effort-based decision-making and did not examine how different types of effort and fatigue interact when making decisions to exert (Hogan *et al*., 2020; Iodice *et al*., 2017; Müller *et al*., 2021).

This study investigated the neural mechanisms by which cognitive fatigue interacts with the brain’s valuation and decision-making circuitry when making choices to exert physical effort. Behaviorally, we hypothesize that cognitive fatigue, induced by repeated working memory exertion, will result in increased feelings of fatigue in both the cognitive and physical domains. This hypothesis is informed by studies that have examined the crosstalk between cognitive fatigue and physical exertion, which found that cognitive fatigue inflated individuals’ perceptions of physical effort (Pageaux, 2014; Pageaux and Lepers, 2016; Harris and Bray, 2019). We hypothesize that fatiguing cognitive exertion will result in an exaggerated subjective valuation of physical effort that manifests as diminished risk preferences for prospective physical effort. When individuals are faced with exerting a certain amount of physical effort versus a risky option involving either a greater amount of effort or no effort, they will be less willing to choose the risky option while in a cognitively fatigued state (compared to a rested state). Our predictions regarding decisions when in a cognitively fatigued state are influenced by studies of physical fatigue and decision-making, which found that increased fatigue was associated with increased subjective valuation and risk preferences for effort (Hogan *et al*., 2020). These behavioral results would suggest a general fatigue signal influencing feelings of effort and choices to exert across effort domains. Neurally, we hypothesize that decisions about prospective effort exertion have their basis in a value signal encoded in the ACC and insula and that the cognitive fatigue state will modulate this value signal. Recent studies of physical fatigue and physical effort-based decision-making found that the insular cortex encodes feelings of effort during bouts of exertion and rest and is sensitive to changes in effort value as a function of physical fatigue (Hogan *et al*., 2020; Meyniel *et al*., 2013; Meyniel *et al*., 2014). We hypothesize that brain regions specifically responsible for executing cognitive effort will be functionally coupled with effort valuation regions such as the insula and that this network will inform effort-based decision-making when in a fatigued state. Together, these hypotheses form an account of how different types of effort and fatigue interact at the levels of brain and behavior to influence effort-based choice.

## Results

To study how decisions about physical effort are influenced by cognitive fatigue, we scanned participants’ brains with functional magnetic resonance imaging (fMRI) while they made risky choices about prospective physical effort before and interspersed with bouts of fatiguing cognitive exertion. The first session of choices was used to characterize participant-specific subjective valuations of physical effort in a baseline, rested state (Figure 1A). After this baseline choice phase, participants performed blocks of cognitive exertion trials in the form of an n-back working memory task (Figure 1B). Participants alternated between blocks of physical effort choice trials and fatiguing cognitive exertion trials (Figure 1D) and rated their cognitive and physical fatigue levels after exertion (Figure 1C). The blocks of exertion trials were meant to maintain participants in a cognitively fatigued state and minimize the possibility of recovery during choice. All the choices were for prospective effort, and at the end of the experiment, ten trials were randomly selected to be played out so that participants’ decisions had actual consequences.

**Figure 1.**
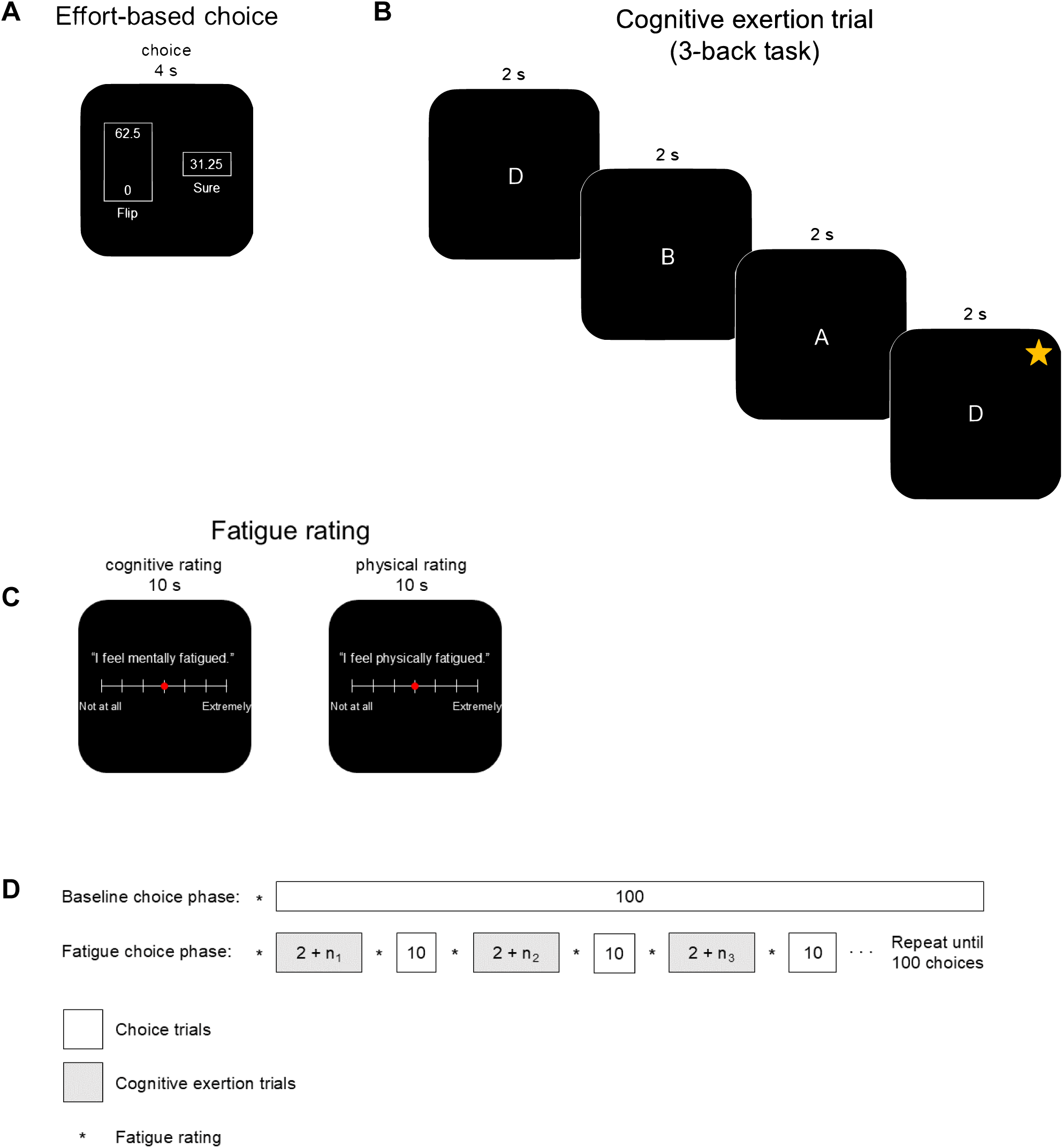
Experimental paradigm. **(A)** During effort-based choice trials, participants completed a series of choices involving the selection of one of two options: a risky option to exert a large amount of physical effort or no effort with equal probability (“Flip”) or exerting a lower amount of physical effort with certainty (“Sure”). Effort amounts were presented on a 0 to 100 scale, which participants were trained on during an association phase before making effort-based choices. An effort level of zero corresponded to no physical exertion and 100 to 80% of a participant’s maximum exertion. To study the effects of cognitive fatigue on effort-based decision-making, blocks of cognitive exertion trials were interspersed with blocks of effort-based choice. **(B)** Cognitive fatigue was induced by having participants perform repeated 3-back working memory trials. Participants were instructed to track a sequence of pseudorandomized letters and identify whether the current letter onscreen (starred “D”) matched the letter appearing three frames previously. **(C)** Participants were queried about their feelings of cognitive and physical fatigue between blocks of physical effort choices and fatiguing cognitive exertion. **(D)** Experiment schedule. The experiment comprised a baseline choice phase followed by a fatigue choice phase, both performed while participants were scanned with fMRI. Participants were questioned about their cognitive and physical fatigue ratings at the beginning and end of each choice block. The baseline choice phase, designed to assess effort preferences in a rested state, comprised 100 randomly presented choices to exert prospective physical effort. In the fatigue choice phase, the same 100 physical effort choices were distributed into 10-trial choice blocks interspersed with blocks of effortful cognitive exertion. During cognitive exertion blocks, participants performed 3-back working memory trials until two sequences were successfully completed (*n_i_* indicates the additional number of 3-back tasks before participants reached two successful completions). This process continued until ten back-to-back blocks of cognitive exertion and choice tasks had been completed.

Before performing fatiguing cognitive exertions, the majority of participants exhibited *ρ_baseline_* > 1, indicating increasing sensitivity to changes in subjective physical effort cost as objective effort level increases (mean *ρ_baseline_* = 2.25 (SD = 1.34); two-tailed one-sample t-test against the null hypothesis that *ρ_baseline_* = 1: *t*_25_ = 4.78, *p* ≪ 0.001). *ρ_baseline_* > 1 corresponds to participants being risk averse for effort. As in our previous work, there was considerable individual variability in participants’ *ρ_baseline_*, reflecting individual differences in baseline subjective preferences for effort (Hogan *et al*., 2019; Hogan *et al*., 2020; Umesh *et al*., 2020).

### Repeated cognitive exertion results in fatigue

Participants’ ratings of cognitive and physical fatigue increased through repeated cognitive exertion (Figure 2A). Ratings of cognitive fatigue significantly increased between the baseline and first session of the fatigue choice phase (average change in cognitive fatigue rating: 1.03 SD; two-tailed paired-sample t-test: *t*_25_ = 3.45, *p* < 0.01), and there was a trend of increased fatigue ratings with progressive blocks of cognitive exertion (hierarchical linear model: *β* = 0.23, *t*_518_ = 12.44, *p* ≪ 0.001). While physical fatigue ratings did not significantly increase between the baseline choice phase and the first session of the fatigue phase (average change in physical fatigue rating: −0.09 SD; two-tailed paired-sample t-test: *t*_25_ = −0.25, *p* = 0.81), there was a trend of increased ratings of physical fatigue with progressive blocks of cognitive exertion (hierarchical linear model: *β* = 0.14, *t*_518_ = 3.74 *p* < 0.01). The rate at which cognitive fatigue ratings increased over progressive exertion blocks was significantly greater than that for physical fatigue ratings (average difference in the slope of cognitive and physical fatigue ratings: 0.10 SD/block; two-tailed paired-sample t-test: *t*_25_ = 2.70, *p* < 0.05), and the rate at which participants ratings of cognitive and physical fatigue increased over exertion blocks was significantly correlated (Figure 2B; Spearman’s *ρ* = 0.42, *p* < 0.05) – individuals with greater rates of increase in cognitive fatigue also had higher rates of increase in physical fatigue. These results suggest that cognitively fatiguing exertion increases feelings of fatigue in both the domains of cognitive and physical effort and are consistent with a general feeling of fatigue that pervades across the different types of effort.

**Figure 2.**
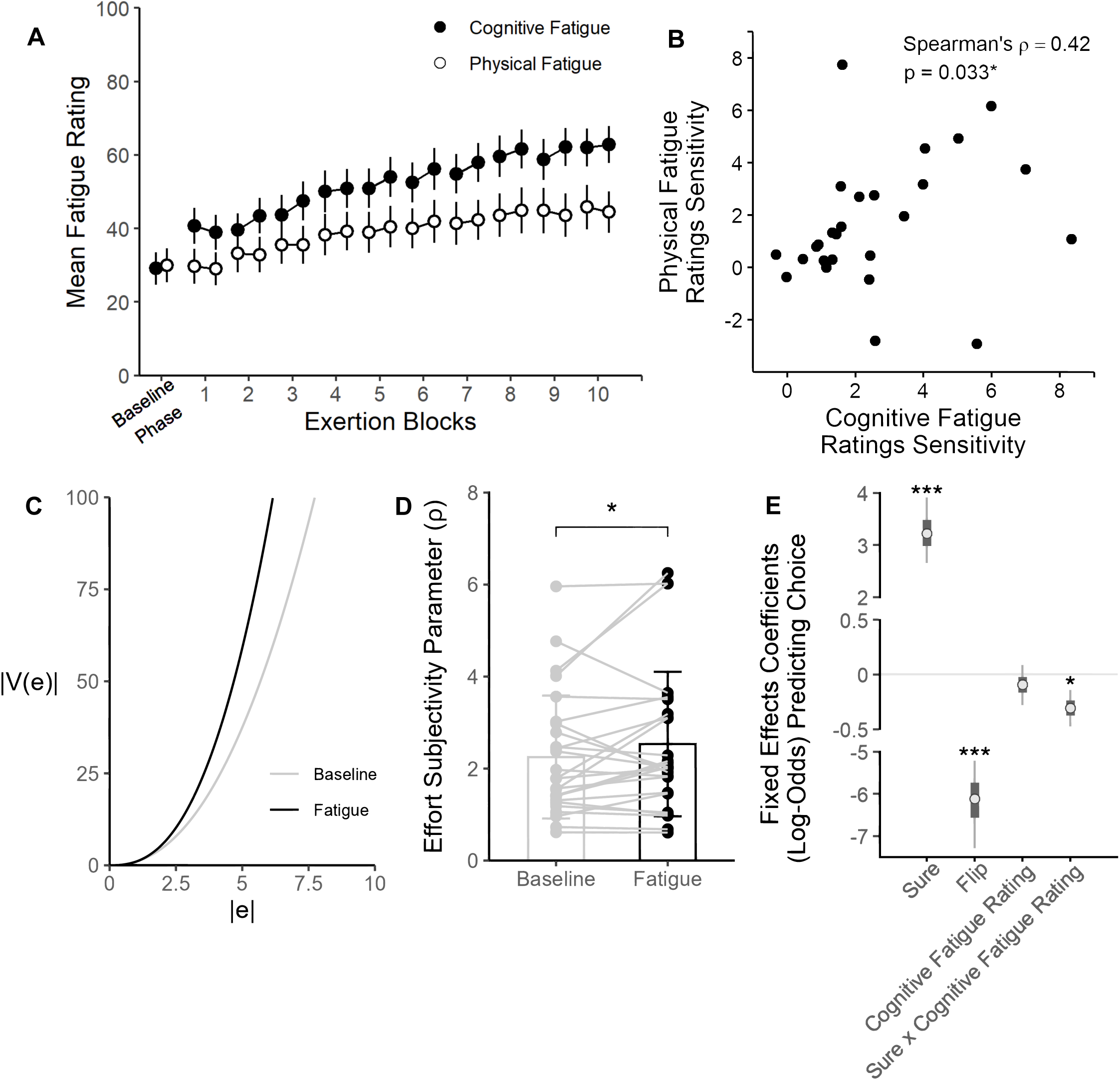
Behavioral results (n = 25). **(A)** Self-reported group-mean cognitive and physical fatigue ratings. Baseline fatigue ratings were collected at the start of the baseline choice phase, and all subsequent ratings were collected before and after each working memory block in the fatigue choice phase. Lines connecting points indicate ratings from the same cognitive fatigue block. Both cognitive and physical fatigue increased significantly throughout the fatigue choice phase; however, cognitive fatigue ratings increased at a greater rate than physical fatigue ratings (average difference in the slope between cognitive and physical fatigue ratings: 0.10 SD/block; two-tailed paired-sample t-test: *p* < 0.05). Error bars indicate SEM. **(B)** Participants’ sensitivity to increasing cognitive and physical fatigue ratings were positively correlated. Participants who reported more rapid increases in cognitive fatigue also reported greater increases in physical fatigue. **(C)** The function used to model the subjective cost of physical effort. This function takes the form of *V*(*x*) = −(−*x*)*^ρ^*. Effort cost functions using mean values of the ρ estimates are indicated by the solid lines (baseline: gray; fatigue: black). Undergoing fatiguing exertions increases the marginal cost of effort. To better illustrate the cost functions, the x- and y-axes shown are not to the same scale. **(D)** The effort subjectivity parameter (ρ) increased significantly between the baseline and fatigue choice phases. A significant increase in ρ indicates that, compared to baseline, exertion-induced fatigue makes the subjective value of physical effort even more costly to participants. Error bars indicate SEM. One-tailed paired-sample t-test: \**p* < 0.05. **(E)** Bayesian hierarchical logistic regression predicting choices to select the risky option during the fatigue choice phase. An interaction between cognitive fatigue rating and the value of the sure option increases the likelihood of individuals selecting the sure option. The asterisks show significant regressors (*: p < 0.05; **: p < 0.01; ***: p < 0.001). Bars indicate standard deviations, and lines are 95% credible intervals of the posterior distributions of each parameter.

Perceptions of fatigue can be influenced by objective decreases in task performance, an effect called performance fatiguability (Kluger *et al*., 2013). Performance fatiguability may manifest as decreased reaction time or task success rate. To evaluate if performance fatiguability may contribute to participants’ fatigue ratings, we evaluated participants’ reaction times and success rates during the progressive blocks of cognitive exertion. We found that participants exhibited lower reaction times (average decrease in RT: −0.017 seconds/block; two-tailed one-sample t-test: *t*_25_ = −5.61, *p* ≪ 0.001) and higher success rates (average percent increase correct: 0.33 %/block; one-tailed one-sample t-test: *t*_25_ = 1.93, *p* < 0.05) over progressive blocks of the n-back cognitive exertion task, revealing patterns of performance that do not align with a fatiguability account. These results suggest that participants experienced increased cognitive and physical fatigue due to time spent on the cognitively fatiguing working memory task rather than performance changes in the task.

### Cognitive fatigue-induced changes in physical effort value

Compared to the baseline choice phase, participants were more risk averse for physical effort during the cognitive fatigue choice phase. Most participants were less willing to take the chance of having to exert large amounts of physical effort, suggesting that their sensitivity to marginal changes in physical effort cost increased while in a cognitively fatigued state (Figure 2C shows group-averaged costs functions for physical effort for the baseline and fatigue choice phases). These cognitive fatigue-induced increases in physical effort cost and risk preferences manifested as a significant increase in *ρ_fatigue_* compared to *ρ_baseline_* (Figure 2D; mean Δ*ρ* = 0.28 (*SD* = 0.82); one-tailed paired-sample t-test: *t*_25_ = 1.76, *p* < 0.05). The parameter *τ*, which represents participants’ randomness in choice, was not significantly different between the baseline and fatigue choice phases (Δ*τ* = 0.13 (*SD* = 0.64); two-tailed paired-sample t-test: *t*_25_ = 0.90, *p* = 0.38), indicating that increased fatigue did not have a significant effect on the variability in a participant’s choices when comparing between rested and cognitively fatigued states.

To capture how cognitive fatigue influences effort-based choices over the course of repeated cognitive exertion, we designed a series of Bayesian hierarchical logistic regression models to measure the effects of cognitive and physical fatigue on the propensity to choose the risky physical effort option over the fatigue phase. We found that an interaction between cognitive fatigue rating and the offered sure value had a significant effect on choice behavior (Figure 2E; Bayesian hierarchical logistic regression: *β* = −0.31, *SE* = 0.10, 95% *CI* = [−0.52, −0.11], *R* = 1.00, *ESS* = 4,490; see Supplementary Figure 2 and Tables 1, 2 for quality analysis of Bayesian modeling), indicating that as cognitive fatigue increased, participants’ willingness to choose the sure option over the risky option increased. In a similar model using physical fatigue as a predictor of choice, we did not find a significant relationship between physical fatigue ratings and the value of the sure option (Supplementary Figure 1), suggesting that, although participants experienced increasing fatigue in both domains, only cognitive fatigue had a significant effect on choice behavior regarding physical effort exertion. A model comparison showed that the model that included cognitive fatigue ratings better described choice behavior than the physical fatigue rating model (cognitive fatigue model: *WAIC* = 900.0; physical fatigue model: *WAIC* = 912.8).

### Neural encoding of physical effort value

We found several brain regions, including the dorsal anterior cingulate cortex and bilateral insula, were sensitive to the difference between chosen and unchosen physical effort value across the baseline and fatigue choice phases (Figure 3A). Brain activity in these areas increased for the chosen effort option compared to the unchosen option, across both the baseline and fatigue choice phases. This finding is consistent with previous studies of effort-based decision-making that have identified these regions as being implicated in effort valuation (Chong *et al*., 2017; Hogan *et al*., 2019; Hogan *et al*., 2020; Klein-Flügge *et al*., 2016; Meyniel *et al*., 2013; Meyniel *et al*., 2014).

**Figure 3.**
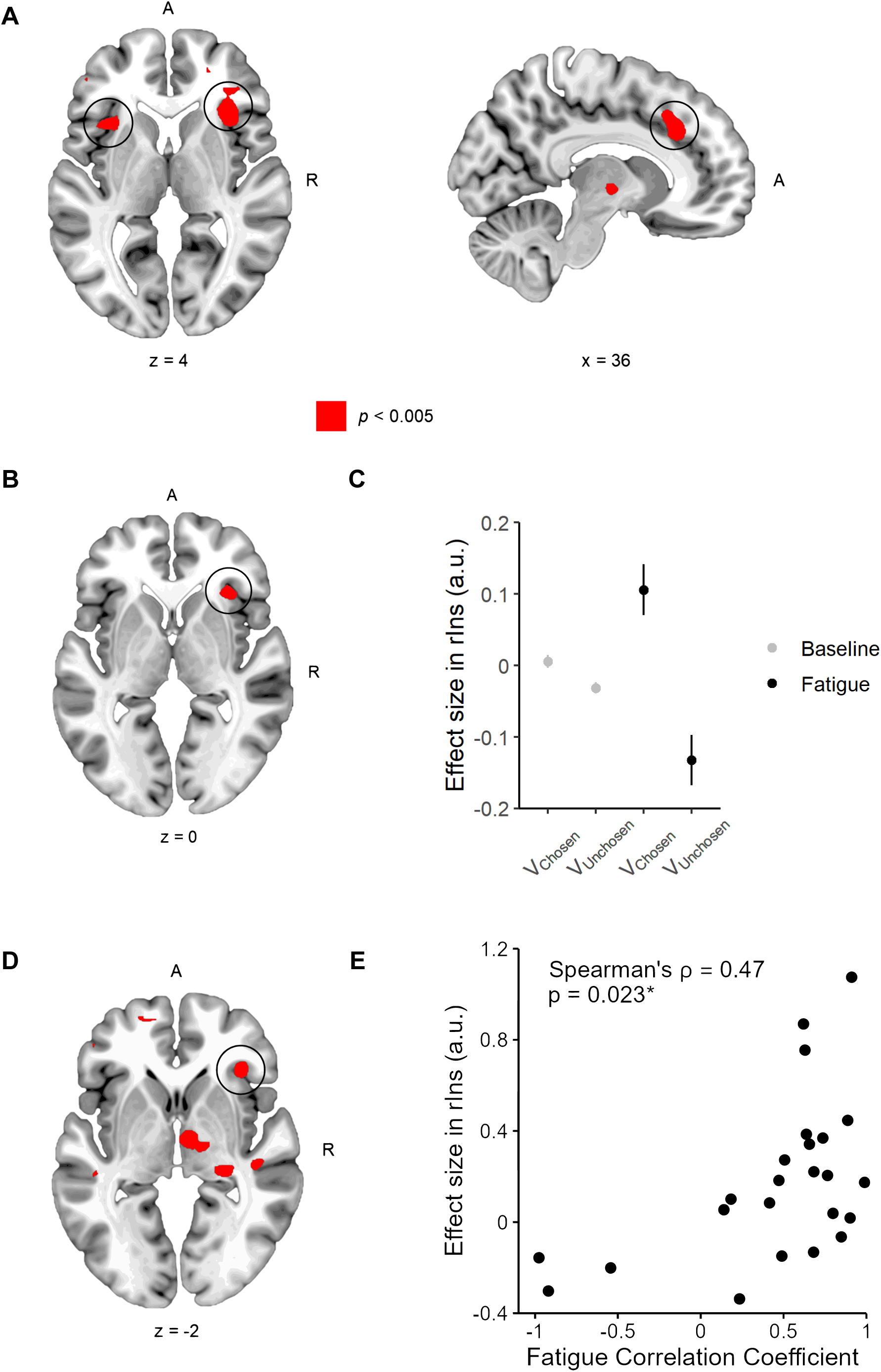
Neural representations of physical effort value (n = 24). **(A)** General physical effort value encoding. Whole brain results thresholded at voxelwise p < 0.005. Activity in bilateral insula (rIns: peak = [34, 24, 4]; lIns: peak = [-38, 18, 2]; small volume corrected p < 0.05 in a priori ROI) illustrates the difference between chosen and unchosen effort value at the time of choice, across the baseline and fatigue choice phases. Activity was also observed in ACC (MNI coordinate: peak = [10, 28, 32]); however, it does not survive small volume correction in our a priori ACC ROI. **(B)** Activity encoding effort value in rIns increases with fatigue. Increased activation in rIns (peak = [34, 26, 0]; small volume corrected p < 0.05 in a priori ROI) indicates the difference of chosen and unchosen effort value between the baseline and fatigue choice phases. **(C)** Effects in rIns (5-mm sphere centered at [34, 26, 0]) for chosen and unchosen effort value between the baseline and fatigue choice phases. This plot was not used for statistical inference (which was carried out in the SPM framework) and is shown to illustrate the pattern of the BOLD signal. Error bars indicate SEM. **(D)** Between participant regression analysis considering the correlation between the progression of cognitive and physical fatigue ratings (Figure 2A), as a covariate for fatigue-induced changes in effort value (peak = [x, y, z]; small volume corrected p < 0.05 in a priori ROI). **(E)** Participants with stronger correlations between physical and cognitive fatigue ratings, rIns exhibited greater sensitivity to changes in physical effort value while in a state of cognitive fatigue. This plot was included to illustrate the relationship between behavior and brain activity and was not used for statistical inference, which was carried out in the SPM framework.

To test for regions of the brain that were sensitive to changes in physical effort value induced by cognitive fatigue, we contrasted the difference between the chosen and unchosen options between the baseline and fatigue choice phases. We found that right anterior insula (rIns) activity was modulated by physical effort value at the time of choice (Figure 3B) and was insensitive to chosen and unchosen effort value in the baseline choice phase (Figure 3C), suggesting that activity in rIns is sensitive to changes in physical effort value resulting from cognitive fatigue. These results align with previous studies of effortful exertion that have suggested that the rIns encodes representations of bodily state that influence decisions regarding bouts of exertion and rest (Meyniel *et al*., 2013; Meyniel *et al*., 2014). Moreover, the region of rIns identified overlaps with the area we previously found for physical effort-based decision-making during physical fatigue (Hogan *et al*., 2020), suggesting that rIns may track the value of physical effort as well as fatigue-induced changes in effort value, regardless of the source of fatigue (i.e., both physical and cognitive fatigue).

To further test how rIns activity at the time of effort choice is modulated by general fatigue, we obtained an independent measure of the associations between participants’ ratings of cognitive and physical fatigue and used it as a covariate in the contrast comparing the baseline and fatigue choice conditions (Figure 3B). The general fatigue measure was obtained by correlating each participant’s increases in cognitive and physical fatigue ratings over the course of repeated cognitive exertion blocks – larger values correspond to a greater agreement between the cognitive and physical fatigue ratings and, thus, greater crosstalk between these fatigue modalities. We found that individuals’ general fatigue metric was significantly related to rIns activity, at the time of choice (Figure 3D, E). Thus, participants with stronger relationships between cognitive and physical fatigue ratings exhibited a higher sensitivity in rIns to fatigue-induced changes in effort value. These results further support the idea of a general fatigue signal for cognitive and physical effort that influences effort-based decisions.

### Increased cognitive fatigue influences physical effort valuation

Next, we evaluated the relationship between cognitive fatigue induced by the working memory task and effort-based decision-making. We reasoned that to make informed decisions about effort, given feelings of fatigue, the brain should incorporate information about the cognitive state (induced by fatiguing cognitive exertion) at the time of choice. To test this idea, we first examined brain areas encoding increased cognitive exertion over the course of the fatigue choice phase. We found that activity in right dorsolateral prefrontal cortex (rdlPFC) increased through repeated cognitive exertion (Figure 4A, B), consistent with previous neuroimaging studies of working memory that have shown this brain region to be related to increased working memory load (Barbey *et al*., 2013; Chong *et al*., 2017; Westbrook and Braver, 2015; Westbrook *et al*., 2019).

**Figure 4.**
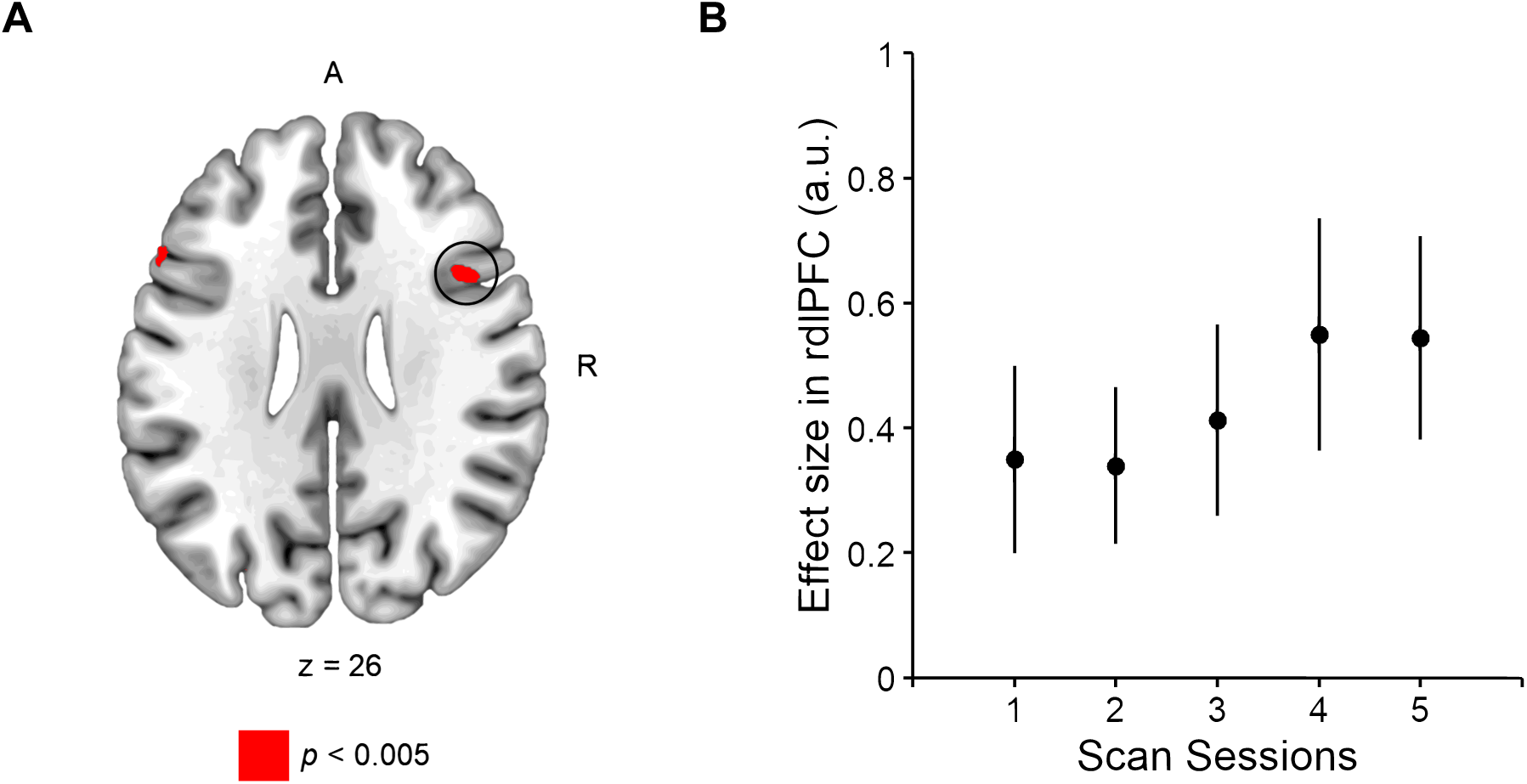
Neural representations of cognitive exertion (n = 24). **(A)** Cognitive exertion-induced changes in brain activity. Whole brain results thresholded at voxelwise p < 0.005. Activity in right dlPFC (peak = [46, 14, 28]; small volume corrected p < 0.05 in a priori ROI) increased with repeated working memory exertion. Activity was also observed in left dlPFC (MNI coordinate: peak = [-56, 20, 22]); however, it does not survive small volume correction in our a priori dlPFC ROI. **(B)** Effects in rdlPFC (5-mm sphere centered at [46, 14, 28]) were positively correlated with exertion block number during the fatigue choice phase. This plot was not used for statistical inference, which was carried out in the SPM framework. Error bars indicate SEM.

Finally, given our hypothesis that information about one’s cognitive fatigue state is incorporated into decisions about physical effort, we tested the idea that the neural circuit modulating effort value representations in rIns might be influenced by computations about cognitively fatiguing working memory instantiated in rdlPFC during choice. To test this hypothesis, we conducted a psychophysiological interaction (PPI) analysis between rIns (seed) and rdlPFC (target) at the time of choice, with baseline/fatigue state as a psychological variable (Figure 5A). This analysis revealed a modulation of functional connectivity between the rIns and rdlPFC as a function of fatigue state, and connectivity was increased in the fatigue choice phase compared to baseline (Figure 5B; mean increase in effect size in rdlPFC: 1.83 a.u.; two-tailed paired-sample t-test: *t*_24_ = 3.03, *p* < 0.01). This analysis provides support for the hypothesis that activity in rdlPFC and rIns are functionally related during effort-based decision-making and suggests that interactions between these brain regions could facilitate the transfer of information about cognitive exertion and fatigue that is used to subserve choices about prospective physical effort.

**Figure 5.**
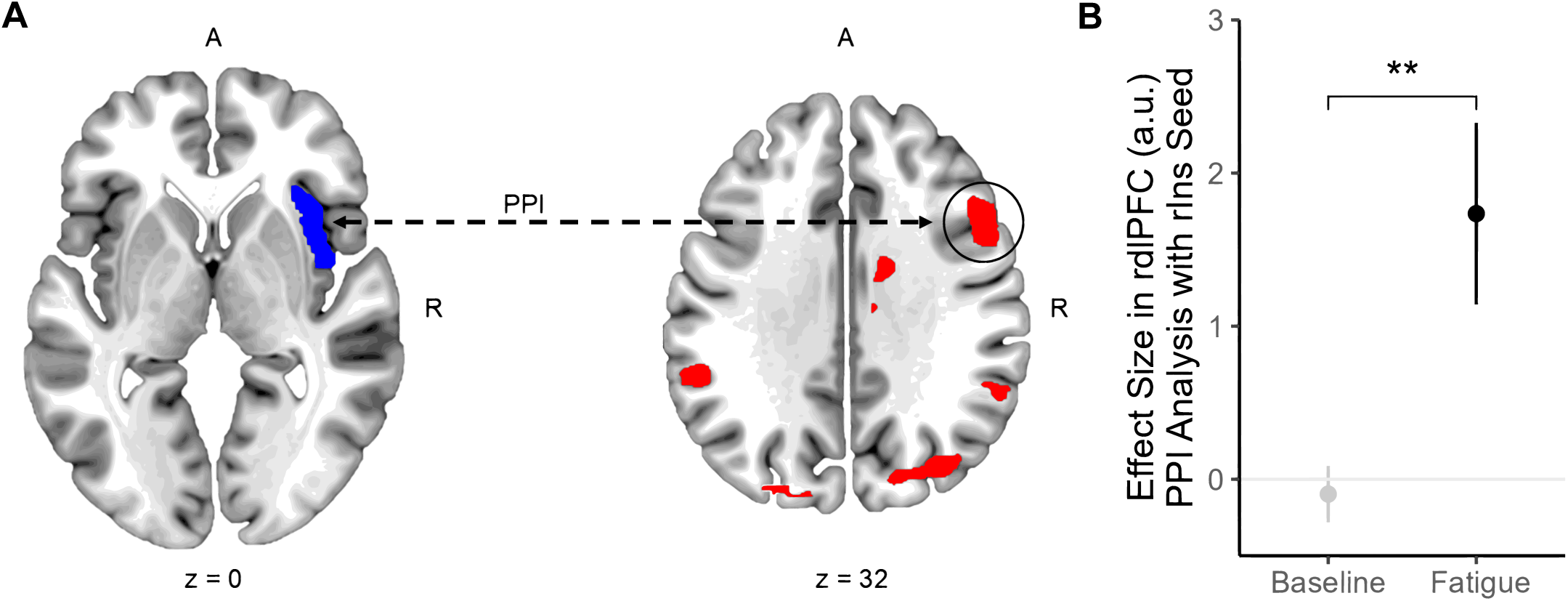
Functional connectivity between rIns and rdLPFC (n = 24). **(A)** Illustration of the psychophysiological interaction (PPI) analysis. We computed a PPI between rIns and rdlPFC with the psychological variable of the baseline/choice phase at the time of choice. **(B)** Effect size in rdlPFC (5-mm sphere centered at [48, 10, 28]) extracted from an a prior ROI showing a modulation in connectivity between this rdlPFC and rIns as a function of fatigue state. Functional connectivity was increased in the fatigue choice phase compared to baseline (average increase in effect size in rdlPFC: 1.83 a.u.; two-tailed paired-sample t-test: *p* < 0.01). Error bars indicate SEM.

## Discussion

We show that repeated cognitive exertion increases feelings of cognitive and physical fatigue and the subjective cost of physical effort. These findings suggest a general fatigue signal influencing behavior across different effort domains. Our neural results reveal that cognitive fatigue-induced changes in physical effort valuation are encoded by rIns, and the functional connectivity between rIns and cognitive exertion-related signals in dlPFC are influenced by fatigue state. These findings are consistent with previous studies that have implicated the anterior insula in an effort valuation network and show that it is sensitive to fatigue-induced changes in effort value (Aridan *et al*., 2019; Chong *et al*., 2017; Hogan *et al*., 2020; Lopez-Gamundi *et al*., 2021; Massar *et al*., 2018; Müller and Apps, 2019; Pessiglione *et al*., 2018; Westbrook and Braver, 2015). However, our results go beyond previous studies by showing that fatigue in one domain of exertion (i.e., cognitive) influences brain signals related to effort valuation in a separate exertion domain (i.e., physical). Our results illustrate a network of brain activity through which disparate effort domains interact to influence decisions to exert.

Effort domain-specific signals are critical for signaling fatigue. In the context of physical effort, fatigue could be related to exertion-induced changes in muscle physiology or motor cortical state (Hogan *et al*., 2020; Müller and Apps, 2019), while it has been suggested that neurotransmitter concentrations in cognitive exertion-related regions are associated with cognitive fatigue (Dobryakova *et al*., 2013; Kok, 2022; McMorris, 2018). Here we show that being in a cognitively fatigued state impacts ratings of physical fatigue and decisions to exert physical effort, suggesting a general fatigue signal that impacts decisions across cognitive and physical domains. We find that a region of rIns that we previously found to be sensitive to cognitive and physical effort-based decision-making while fatigued in those respective domains (Hogan *et al*., 2020; Steward and Chib, 2024; Westbrook and Braver, 2015), also mediates decisions to exert physical effort while in a cognitively fatigued state. At the time of choice, we find that specific working memory-related cognitive exertion signals in dlPFC are functionally coupled to this region of rIns, suggesting that information about task-related neural activity plays a role in effort-based choice. However, our data is not able to distinguish how signals related to cognitive and physical fatigue might be synthesized into a general fatigue signal that underlies choice. rIns is a candidate region that is sensitive to effort decisions in both cognitive and physical fatigue (Chong *et al*., 2017; Müller and Apps, 2019); however, it is not clear if other brain regions encode a general fatigue state across choices and exertion.

It is important to monitor one’s internal state to make decisions about exertion while fatigued. The region of rIns that we have identified as being sensitive to physical effort value while in a cognitively fatigued state has also been shown to be sensitive to cognitive and physical fatigue while making effort choices in those domains of exertion (in which there was no crosstalk between types of effort; Hogan *et al*., 2020; Steward and Chib, 2024). This region of rIns overlaps with the region identified in the computation of interoceptive sense (Craig, 2003; Craig, 2009; Critchley *et al*., 2004). One interpretation of rIns being sensitive to fatigue-induced changes in effort value could be that this region may be required to access effort domain-specific interoceptive feelings, which in turn, influence valuations and judgments of effort. In this framework, rIns could serve as a domain-general node in fatigue judgements. While our study did not directly assess participants’ interoceptive sense related to feelings of cognitive fatigue, it will be important in the future to design experimental paradigms that measure an individual’s interoceptive awareness of cognitive state while also requiring them to make decisions about prospective cognitive and physical exertion. Such an experimental design could allow for the dissociation of interoceptive signals and effort valuation in rIns.

Motivation is another key driver of effortful behavior generally impacted by fatigue. Cognitive and physical fatigue alter the cost-benefit analysis underlying decision-making, where the perceived effort required for tasks diminishes the subjective value of potential rewards, thereby reducing their motivational salience (Chong *et al*., 2017; Iodice *et al*., 2017; Klein-Flügge *et al*., 2016; Massar *et al*., 2018; Westbrook *et al*., 2013). When motivation is low, the effort needed to achieve a reward can seem disproportionately burdensome, making the reward less appealing than it would be in a more motivated state. As fatigue accrues from sustained cognitive or physical exertion, exertion-related neural signals may influence the general brain regions integral to motivated behavior, such as the basal ganglia and prefrontal cortex. These signals could modulate internal assessments of whether future rewards justify the required effort. Our study examined how cognitive fatigue shapes decisions involving physical effort, revealing a functional network, including the dlPFC and rIns, which may be critical in motivating choices to exert effort. These regions potentially mediate the interplay between subjective effort valuation and motivated decision-making under fatigue. While our experiment did not test the influence of incentive motivation on decisions to exert, instead focusing on effort valuation in isolation, reward motivation would likely have a general impact on fatigue that influences decisions across effort domains.

Through a combination of behavioral and neural analysis, we show that cognitive fatigue impacts feelings of physical fatigue and decisions to exert physical effort. These findings suggest a domain-general fatigue network that draws on exertion-related neural signals to influence judgments of cognitive and physical effort. We show a mechanism by which representations of physical effort value in rIns are modulated by cognitive fatigue-induced changes in rdlPFC, and that these brain regions are functionally connected as part of an effort-fatigue network that influences effort-based decision-making.

## ACKNOWLEDGEMENTS

This work was supported by the Eunice Kennedy Shriver National Institute of Child Health & Human Development of the National Institutes of Health under Award Number R01MH119086 and the National Institutes of Mental Health R01HD097619.

## Methods

### Experimental setup

The presentation of visual stimuli and acquisition of behavioral data was accomplished using custom PsychoPy scripts (Pierce *et al*., 2019). During fMRI, visual feedback was presented by a projector located at the back of the room. Participants viewed a reflection of the projection via a mirror attached to the scanner head coil.

A hand-clench dynamometer (TSD121B–MRI, BIOPAC Systems, Inc., Goleta, CA) recorded grip force exertion. Signals from this sensor were sent to our custom-designed software for real-time visual feedback of participants’ exertions. Participants were instructed to exert a grip force on the sensor in their dominant hand while comfortably holding their arm to the side.

We used an MRI-compatible multiple button-press response box (Cedrus RB-830, Cedrus Corp., San Pedro, CA) held in the left hand to record participant decisions while in the scanner.

### Participants

All participants were right-handed and prescreened to exclude any individuals with a history of neuropsychiatric conditions. The Johns Hopkins School of Medicine Institutional Review Board approved this study, and all participants provided informed consent.

A total of 38 healthy participants were recruited from the Johns Hopkins community. Of these, 13 participants were excluded from behavioral analyses, and 14 were excluded from neuroimaging analyses for one or more reasons. First, participants were excluded if they did not complete the study due to complications during scanning (n = 4). Second, participants were excluded if their choice parameters (*ρ* and *τ*) were outliers (> 2 standard deviations from the mean; n = 2). Third, participants were excluded if their cognitive ratings displayed no variance (i.e., they did not increase or decrease throughout the experiment; n = 3). Fourth, participants were excluded if they made nonsensical choices (n = 4). Finally, one participant was included in the behavioral analysis but excluded from neuroimaging analysis due to excessive head movement. The final analysis included N = 25 participants (N = 24 for neuroimaging analysis) in total (mean age ± standard deviation, 24 ± 5y; range, 18 – 39y; 11 males)

### Experimental paradigm

Before the experiment, participants were informed they would receive a fixed show-up fee of $50. They were told that this fee did not depend on their performance or behavior during the experiment. The association, assessment, and choice phases of the experiment described below are similar to those we have previously used (Culbreth *et al*., 2024; Hogan et al., 2019; Hogan et al., 2020; Hu *et al*., 2022; Padmanabhan *et al*., 2023; Umesh *et al*., 2020).

The experiment began by measuring participants’ maximum voluntary contraction (MVC), by selecting the maximum grip exertion on the hand-clench dynamometer over three successive trials. During these exertions, participants did not have knowledge about subsequent phases and were encouraged to squeeze with their maximum force.

Following the MVC phase, participants underwent an association phase during which they learned to associate effort levels (relative to MVC) with the corresponding force they exerted on the dynamometer (Supplementary Figure 3A). Effort levels were presented on a scale ranging from 0 effort units (no exertion) to 100 effort units (80% of a participant’s MVC). Participants proceeded through a randomized order of training blocks, each consisting of five training trials for a single target effort level, ranging from 10–80 effort units in increments of 10. We did not implement association trials at the highest levels of exertion (i.e., 100% of a participant’s MVC) to minimize the risk of participants becoming physically fatigued during this phase. Each trial of a training block began with the numeric presentation of the target effort level (2 s), followed by effort exertion with visual feedback in the form of a black vertical bar, similar in design to a thermometer, which increased in level the harder participants gripped the dynamometer (4 s). The bottom and top of this effort gauge represented effort levels 0 and 100, respectively. Participants were instructed to reach the target zone (±5 effort units of the target) as fast as possible and maintain their force within the target zone for as long as possible for 4 s. Visual indication of the target zone was colored green if the effort produced was within the target zone, and red otherwise. After exertion, if participants were within the target zone for more than two-thirds of the trial time (2.67 s), the trial was a success. Participants were provided feedback regarding their success or failure at maintaining the target effort after each trial. To minimize participants’ fatigue, a fixation cross (2–5 s) separated the trails within a training block, and 60 s of rest were provided between training blocks.

Following the association phase, participants performed an assessment phase, during which they performed an effort recall task that gauged their understanding of the association between the effort levels and their physical exertion (Supplementary Figure 3B). All the effort levels from the association phase (10–80 in increments of 10 effort units) were presented randomly six times each. Each assessment trial began with the display of a black horizontal bar that participants were instructed to fill by grip exertion on the dynamometer. Visual feedback turned red to green once the target effort level was reached. A full bar did not correspond to an effort level of 100 as in the previous phase; here, it represented the target effort level required on each trial. Participants were told to reach the target zone as fast as possible, maintain their force production as long as possible, and estimate their effort level during exertion (4 s). Following this exertion, participants were presented with a number line ranging from 0 to 100 and told to select the effort level they believed they had just exerted. Selection was achieved by moving the computer mouse to the rating and clicking the left mouse button to finalize the response. Participants had 4 s to make this effort assessment; if they failed, the trial was counted as missed. No feedback was given to participants as to the accuracy of their selection. After each selection, a fixation cross (2–5 s) appeared on the screen to provide a rest period between trials. A longer rest period of 60 s was provided halfway through the phase.

Following the assessment phase, participants were introduced to the n-back task, a cognitive effort paradigm commonly used to engender cognitive exertion through repeated use of working memory (Westbrook et al., 2013). We chose this task because we could operationalize cognitive effort by modulating the working memory load by varying the value of ‘n’ (Westbrook et al., 2013). In this experiment, we employed a 3-back version of the n-back task, wherein participants monitored a sequence of letters and identified any letter (i.e., target) that matched the one shown 3 frames previously (Figure 1B). Participants completed a practice session of the 3-back task consisting of 40 letters, 10 of which were targets. Participants identified target and non-target letters with keyboard presses. Participants were required to complete the practice session (correctly identifying five or more targets) before moving to the main experiment in the scanner.

To investigate the influence of cognitive fatigue on behavioral and neural representations of physical effort valuation, we scanned participants’ brains with fMRI while they made decisions about prospective physical effort. This was done before and after participants performed repeated 3-back tasks and reported their cognitive and physical fatigue levels. Before entering the scanner, participants were told that 10 of their decisions would be randomly selected and carried out at the end of the experiment and that they would have to remain in the testing area until they successfully achieved the selected exertions. Participants were also informed that they should treat each effort decision as separate and independent from the others.

During the scanning portion of the experiment, participants reported their baseline levels of cognitive and physical fatigue via a 7-point Likert scale (10 s), which asked them to indicate their level of agreement (on a scale of “Not at all” to “Extremely”) with the statement “I feel cognitively/physically fatigued” (Figure 1C). The order in which cognitive and physical fatigue questionnaires were presented was randomized. Fatigue levels were reported by pressing a hand-held button box with the left hand’s second, third, and fourth digits. For the remainder of the baseline choice phase, which was designed to gauge effort preferences in a pre-fatigued state, participants were presented with a series of effort choices between two options shown (4 s): a risky decision to exert either a large amount of physical effort or no effort with equal probability (“Flip”), or exerting a small amount of physical effort with certainty (“Sure”) (Figure 1A). (See Table 3 in the Supplementary Materials for the full choice set.) Participants selected between the two options by pressing the same button box with either the third or fourth digits of the left hand. Choices were not realized within the scanner. One hundred effort choices were presented consecutively in a randomized order. Participants were encouraged to make a choice on every trial; however, missing a trial was not penalized. Missed trials (including those for the fatigue surveys) were recorded as such and were not repeated. Previous studies used a similar effort-based decision-making task (Hogan *et al*., 2019; Hogan *et al*., 2020).

Following the baseline choice phase, participants completed the fatigue choice phase of the experiment, in which they alternated between cognitively fatiguing working memory blocks and choice blocks (Figure 1D). A working memory block consisted of fatigue surveys (the same as those used in the baseline choice phase) immediately preceding and following two successful bouts of the 3-back task. Participants completed the 3-back task by pressing the button box with the second or third digits of the left hand. The 3-back task was repeated until participants reached two successful completions. Following a working memory block, participants performed a choice block of 10 effort decisions pseudo-randomly sampled from the same set used in the baseline choice phase.

Following the fatigue choice phase, participants exited the scanner and completed 10 choice trials drawn from decisions made during both the baseline and fatigue choice phases. Participants remained in the testing area until they achieved the target exertions from the chosen trials.

### MRI protocol

A 3 Tesla Philips Ingenia Elition X-series MRI scanner and radio frequency coil was used for all MR scanning sessions. High-resolution structural images were collected using a standard MPRAGE pulse sequence, providing full brain coverage at a resolution of 0.946 mm × 0.946 mm × 1 mm. Functional images were collected at an angle of 30° from the anterior commissure-posterior commissure (AC-PC) axis, which reduced signal dropout in the orbitofrontal cortex (Deichmann et al., 2003). Forty-eight slices were acquired at a resolution of 1.87 mm × 1.88 mm × 3 mm, providing whole brain coverage. An echo-planar imaging (FE EPI) pulse sequence was used (TR = 2800 ms, TE = 30 ms, FOV = 240, flip angle = 70°).

### Effort choice analysis

We used a two-parameter model to capture the subjective cost of effort. We assumed a participant’s cost function *V*(*x*) for physical effort *x* to be of the form:

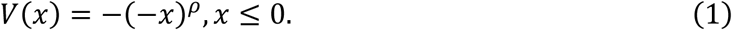

Here, *x* is defined as the objective value of effort and is negative to match our assumption that effort is perceived as a cost. The parameter *ρ* represents sensitivity to changes in subjective effort value as the value of *x* changes. A large *ρ* represents a high sensitivity to increases in objective effort. If *ρ* = 1, then the subjective cost of effort is the objective cost.

Representing the effort levels as prospective costs, and assuming participants combine probabilities and utilities linearly, the relative value between the risky and sure effort options can be written as:

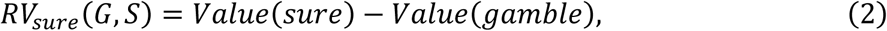

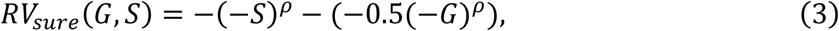

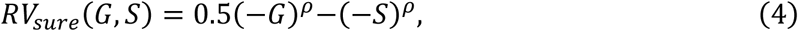

where *RV_sure_* is the difference between the two options, and both *G* < 0 and *S* < 0 for all trials.

We used a softmax function to calculate the probability that a participant chooses the sure option on the *kth* choice trial:

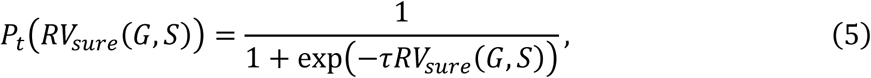

where *τ* is a non-negative temperature parameter measuring the stochasticity of a participants’ choices. If *τ* = 0, then choices were made randomly.

Using maximum likelihood estimation, we extracted the *ρ* and *τ* parameters for each participant, using 100 trials of effort choices. A participant’s choice is denoted by *y ɛ* {0,1}. *y* = 1 indicates the sure option was chosen. Parameters were estimated by maximizing the following likelihood function individually for each participant:

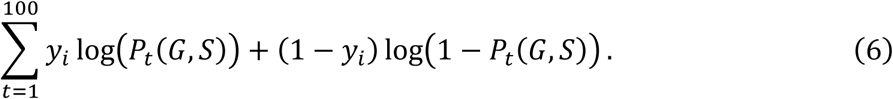

Parameters were estimated separately for the baseline and fatigue choice phases. We acquired *ρ_baseline_*, *τ_baseline_*, *ρ_fatigue_*, and *τ_fatigue_* parameters for each participant.

### Hierarchical modeling of effort choices

We used Bayesian hierarchical logistic regression using the *brms* package (Bürkner, 2017) in R to estimate the trial-to-trial effects of cognitive and physical fatigue on choice behavior. We opted for a Bayesian analysis to account for the quasi-separation in our choice set due to the inclusion of the catch trials in which the raw value of the flip option was always lower than the sure alternative. Such separation can inflate regression coefficients and influence interpretation of the results; thus, we employed penalized regression through the Bayesian method of setting priors for the fixed effects of our model. Before estimating any models, we specified a seed for the pseudo-random number generator; this seed can be downloaded from the Supplementary Materials for exact reproducibility of the model results. We followed the BARG method (Kruschke, 2021) in detailing our model interpretation and reporting.

We estimated the following model to measure the influence of cognitive and physical fatigue on effort choice:

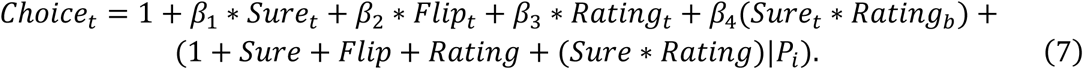

*Choice_t_* is a binary variable representing whether the sure or risky option was picked (0 = sure, 1 = flip) on a given trial *t*, *Sure_t_* is the expected value of the sure option on trial *t*, *Flip_t_* is the expected value of the risky option on trial *t*, *Rating_b_* is the most recent cognitive or physical fatigue rating, and *P_i_* is a categorical identifier for each participant. Given individual differences between participants in their valuations of physical effort and initial and subsequent fatigue levels, maximal models were built with random effects for slope and intercept. All regressors were z-scored before input into the model, and separate models were estimated for cognitive and physical fatigue ratings. Model results for each parameter are reported as the mean, standard deviation, and 95% credible intervals of the posterior distribution. Significant results were identified by observing whether the 95% credible intervals for each parameter crossed 0.

Models were assigned to the Bernoulli statistical family with a logit link function to account for the dual nature of the effort choices. We used broad, weakly informative priors in the form of *normal*(0, 10) for all fixed effects, assuming that the coefficient for the sure option would be positive (indicating that an increasing value of sure option increases the odds of picking the risky option) and that the coefficients for the risky option, fatigue rating, and the interaction between sure and fatigue rating would all be negative (indicating that an increasing value of the risky option and increasing fatigue reduces the odds of picking the flip option). Both models for cognitive and physical fatigue were estimated using the *brms* package’s Markov chain Monte Carlo method, which had 4 chains and 2,000 iterations per chain. The first 1,000 iterations served as the warm-up period, while the remaining iterations acted as the sampling period.

We performed a posterior predictive check on each cognitive and physical fatigue model by qualitatively observing whether the posterior predictive distributions generated by the *pp_check* function in the *brms* package encapsulated the actual distributions (Supplementary Figure 2). Model comparison was conducted by comparing weighted AIC scores generated by the *WAIC* function in the *brms* package.

We confirmed the reliability and efficiency of each model by observing the *R* convergence diagnostic and ESS of each relevant parameter—namely, sure option, risky option, fatigue rating, and sure*fatigue rating. We were satisfied if *R^* < 1.05 and *ESS* > 1,000 for each relevant parameter. To ensure the reliability of our results, we performed a sensitivity analysis by conducting Bayesian hierarchical logistic regression with other broad priors that were more or less informative than the prior described above. In the order of most informative to least informative, these priors included: *normal*(0, 1), *normal*(0, 10^6^), and *uniform*(1, *∞*), the final prior being the *brms* package’s default prior.

The results of this analysis can be viewed in Supplementary Table 1.

### Image processing and fMRI statistical analysis

#### Image preprocessing

The SPM12 software package was used to analyze the MRI data (Wellcome Trust Centre for Neuroimaging, Institute of Neurology; London, UK). A slice-timing correction was applied to the functional images to adjust for different slices within each image being acquired at slightly different time points. Images were corrected for head motion by registering all images to the first image, spatially transformed to match a standard echo-planar imaging template brain, and smoothed using a 3D Gaussian kernel (8 mm FWHM) to account for anatomical differences between participants. Following pre-processing, the data were analyzed statistically with a general linear model (GLM).

#### General linear model

A GLM was used to estimate participant-specific (first-level), voxel-wise, statistical parametric maps (SPMs) from the fMRI data. Our GLM included a categorical boxcar regressor for choice trials, in both the baseline and fatigue choice phases, beginning when a choice was presented and ending when a decision was made. This regressor included unorthogonalized parametric modulators corresponding to the objective value of the risky and sure effort options. Missed choice trials were modeled as a separate nuisance regressor. In the fatigue choice phase, another categorical boxcar regressor was used to model blocks of the working memory (3-back) task, beginning with the first round and ending after the second completed round (unsuccessful rounds were included in this timeframe). Finally, regressors modeling head motion as derived from the affine part of the realignment procedure of the preprocessing pipeline were included in the model.

The regressors included in our imaging model were as follows:

1. Choice trials during the baseline choice phase (Box-car categorical regressor beginning at the time of choice presentation and ending at the time of response)

a. Parametric modulator: Value of the chosen option
b. Parametric modulator: Value of the unchosen option
2. Choice trials during the fatigue choice phase (Box-car categorical regressor beginning at the time of choice presentation and ending at the time of response)

a. Parametric modulator: Value of the chosen option
b. Parametric modulator: Value of the unchosen option
3. Working memory blocks during the fatigue choice phase (Box-car categorical regressor beginning at the time of presentation of the first round of the 3-back task and ending at the conclusion of the second successful round of the 3-back task)
4. Choice trials in which no decision was made in the allotted time (i.e., missed trials)
5. Regressors modeling head motion as derived from the affine part of the realignment procedure of the preprocessing pipeline.

We used these first-level models to create group-level (second-level) models to test for brain areas that were generally sensitive to effort value and cognitive exertion. We created contrasts using the aforementioned parametric modulators for chosen and unchosen effort values, at the time of choice, to identify brain areas sensitive to differences between chosen and unchosen options, both across and between the baseline and fatigue choice phases. To identify brain regions encoding decision values for effort, regardless of fatigue state, we created a contrast that modeled the difference between chosen and unchosen effort value. This contrast was constructed by subtracting the parametric modulator for the unchosen risky and sure options (1.b and 2.b) from the chosen risky and sure options (1.a and 2.a). Additionally, we tested for brain regions encoding decision values that were influenced by the effect of fatigue by taking the difference between the value of the chosen and unchosen options between the fatigue and baseline choice phases ([2.a – 2.b] – [1.a – 1.b]). To test for brain regions sensitive to increases in cognitive fatigue, we created a contrast that assigned linearly increasing weights to the categorical regressor for each n-back working memory block across the duration of the fatigue choice phase.

#### Statistical inference

We analyzed brain signals related to chosen effort value within independent ROIs taken at peak coordinates from Neurosynth.org (Gorgolewski *et al*., 2016) when using the term “effort”: right anterior insula (rIns) MNI coordinates (x, y, z) = [36, 22, 0]; left anterior insula (lIns) MNI coordinates (x, y, z) = [−36, 22, 0]; ACC MNI coordinates (x, y, z) = [0, 14, 46]. Brain regions typically involved in working memory processes include right and left dlPFC and we used Neurosynth.org with the search term “working memory” to obtain independent ROIs for these regions: rdlPFC MNI coordinates (x, y, z) = [48, 10, 28]; ldlPFC MNI coordinates (x, y, z) = [-46, 8, 28].

To display modulations in rIns and rdlPFC activity during the fatigue choice phase, we used SPM12’s *marsbar* (Brett *et al*., 2002) and *rfxplot* (Gläscher, 2009) toolboxes to extract effects sizes. Plots used for statistical inference (Figs. 3D, 5B) were created by extracting BOLD activations using 5-mm spheres centered at the peak coordinates inferred from Neurosynth.org (see above). Otherwise, effect sizes were extracted 5-mm spheres at the peak of activity in our data (Figs. 3C, 4B) – these signals were not statistically independent (Kriegeskorte *et al*., 2009), and these plots were not used for statistical inference and used only for illustrative purposes.

#### Psychophysiological interaction (PPI) analysis

We performed a PPI analysis to assess changes in connectivity between the rIns striatum and dlPFC as a function of fatigue state. PPI is a measure of context-dependent connectivity, which explains the activity of other brain regions in terms of the interaction between responses in a seed region and cognitive processes (Friston *et al*., 1997).

The PPI terms were generated by computing formal interactions between the physiological variable (*Y*) and the psychological variable (*P*). The physiological variable *Y* was the blood-oxygen-level-dependent (BOLD) time courses taken from the participant-specific coordinates of peak activation in an anatomical mask of anterior rIns and deconvolved using a model of a canonical hemodynamic response function. The anatomical mask of rIns was generated using SPM’s Neuromorphometrics Atlas from the area labeled “right anterior insula”. To construct the psychological variable *P*, we contrasted the baseline and fatigue conditions at the time of choice, irrespective of effort value. We generated PPI regressors for the rIns using these physiological and psychological variables.

## References

1. Aridan N, Malecek NJ, Poldrack RA, Schonberg T (2019) Neural correlates of effort-based valuation with prospective choices. Neuroimage 185:446–454.

2. Barbey AK, Koenigs M, Grafman J (2013) Dorsolateral prefrontal contributions to human working memory. Cortex 49:1195–1205.

3. Barr DJ, Levy R, Scheepers C, Tily HJ (2013) Random effects structure for confirmatory hypothesis testing: Keeping it maximal. Journal of Memory and Language 68:255–278.

4. Bates D, Mächler M, Bolker B, Walker S (2015) Fitting linear mixed-effects models using lme4. Journal of Statistical Software 67:1–48.

5. Brett M, Anton J-L, Valabregue R, Poline J-B (2002) Region of interest analysis using an SPM toolbox [abstract] Presented at the 8th International Conference on Functional Mapping of the Human Brain, June 2-6, 2002, Sendai, Japan. Available on CD-ROM in NeuroImage, Vol 16, No 2.

6. Bürkner PC (2017) brms: An R package for Bayesian multilevel models using Stan. Journal of Statistical Software 80:1–28.

7. Chong TT-J, Apps M, Giehl K, Sillence A, Grima LL, Husain M (2017) Neurocomputation mechanisms underlying subjective valuation of effort costs. PLOS Biology 15: e1002598.

8. Craig AD (2003) Interoception: The sense of the physiological condition of the body. Current Opinion in Neurobiology 13:500–505.

9. Craig AD (2009) How do you feel — now? The anterior insula and human awareness. Nature Reviews Neuroscience 10:59–70.

10. Critchley HD, Wiens S, Rotshtein Pia, Öhman A, Dolan RJ (2004) Neural systems supporting interoceptive awareness. Nature Neuroscience 7:189–195.

11. Culbreth AJ, Chib VS, Riaz SS, Manohar SG, Husain M, Waltz JA, Gold JM (2024) Increased sensitivity to effort and perception of effort in people with schizophrenia. Schizophrenia Bulletin sbae162.

12. Deichmann R, Gottfried JA, Hutton C, Turner R (2003) Optimized EPI for fMRI studies of the orbitofrontal cortex. Neuroimage 19:430–431.

13. Dobryakova E, DeLuca J, Genova HM, Wylie GR (2013) Neural correlates of cognitive fatigue: Cortico-striatal circuitry and effort-reward imbalance. Journal of the International Neuropsychological Society 19:849–853.

14. Eddy MD, Hasselquist L, Giles G, Hayes JF, Howe J, Rourke J, Coyne M, O’Donovan M, Batty J, Brunyé TT, Mahoney CR (2015) The effects of load carriage and physical fatigue on cognitive performance. PLOS One 10: e0130817.

15. Friston KJ, Buechel C, Fink GR, Morris J, Rolls E, Dolan RJ (1997) Psychophysiological and modulatory interactions in neuroimaging. Neuroimage 6:218–229.

16. Gläscher L (2009) Visualization of group inference data in functional neuroimaging. Neuroinform 7:73–82.

17. Gorgolewski KJ, Varoquaux G, Rivera G, Schwarz Y, Ghosh SS, Maumet C, Sochat VV, Nichols TE, Poldrack RA, Poline J-B, Yarkoni T, Margulies DS (2015) NeuroVault.org: A web-based repository for collecting and sharing unthresholded statistical maps of the human brain. Neuroinform 9:1–9.

18. Harris S and Bray ST (2019) Effects of mental fatigue on exercise decision-making. Psychology of Sport and Exercise 44:1–8.

19. Hogan PS, Galaro JK, Chib VS (2019) Roles of ventromedial prefrontal cortex and anterior cingulate in subjective valuation of prospective effort. Cerebral Cortex 29:4277–4290.

20. Hogan PS, Chen SX, Teh WW, Chib VS (2020) Neural mechanisms underlying the effects of physical fatigue on effort-based choice. Nature Communications 11:4026–4040.

21. Hu EJ, Casamento-Moran A, Galaro JK, Chan KL, Edden RAE, Puts NAJ, Chib VS (2022) Sensorimotor cortex GABA moderates the relationship between physical exertion and assessments of effort. Journal of Neuroscience 42: 6121–6130.

22. Iodice P, Calluso C, Barca L, Bertollo M, Ripari P, Pezzulo G (2017) Fatigue increases the perception of future effort during decision making. Psychology of Sport and Exercise 33:150–160.

23. Klein-Flügge MC, Kennerley SW, Friston K, Bestmann S (2016) Neural signatures of value comparison in human cingulate cortex during decisions requiring an effort-reward trade-off. Journal of Neuroscience 36:10002–10015.

24. Kok A (2022) Cognitive control, motivation and fatigue: A cognitive neuroscience perspective. Brain and Cognition 160:105880.

25. Kriegeskorte N, Simmons WK, Bellgowan PSF, Baker CI (2009) Circular analysis in systems neuroscience: The dangers of double dipping. Nature Neuroscience 12:535–540.

26. Kruschke JK (2021) Bayesian analysis reporting guidelines. Nature Human Behavior 5:1282–1291.

27. Kuznetsova A, Brockhoff PB, Christensen RHB (2017) lmerTest Package: Tests in linear mixed effects models. Journal of Statistical Software 82:1–26.

28. Lieberman MD and Cunningham WA (2009) Type I and Type II error concerns in fMRI research: Re-balancing the scale. Social, Cognitive, and Affective Neuroscience 4:423–428.

29. Lopez-Gamundi P, Yao Y-W, Chong TT-J, Heekeren HR, Mas-Herrero E, Marco-Pallarés (2021) The neural basis of effort valuation: A meta-analysis of functional magnetic resonance imaging studies. Neuroscience and Biobehavioral Reviews 131:1275–1287.

30. Massar SAA, Casthó A, Van der Linden D (2018) Quantifying the motivational effects of cognitive fatigue through effort-based decision making. Frontiers in Psychology 9:1–5.

31. McMorris T (2018) Cognitive fatigue effects on physical performance: The role of interoception. Sports Medicine 50:1703–1708.

32. Meyniel F, Sergent C, Rigoux L, Daunizeau J, Pessiglione M (2013) Neurocomputational account of how the human brain decides when to have a break. PNAS 110:2641–2646.

33. Meyniel F, Safra L, Rigoux L, Pessiglione M (2014) How the brain decides when to work and when to rest: Dissociation of implicit-reactive from explicit-predictive computational processes. PLoS Computational Biology 10:e1003584.

34. Moore RD, Romine MW, O’connor PJ, Tomporowski PD (2012) The influence of exercise-induced fatigue on cognitive function. Journal of Sports Sciences 30:841–850.

35. Müller T and Apps MAJ (2019) Motivational fatigue: A neurocognitive framework for the impact of effortful exertion on subsequent motivation. Neuropsychologia 123:141–151.

36. Müller T, Klein-Flügge, Manohar SG, Husain M, Apps MAJ (2021) Neural and computational mechanisms of momentary fatigue and persistence in effort-based choice. Nature Communications 12:4593–4606.

37. Padmanabhan P, Casamento-Moran A, Kim A, Gonzalez AJ, Pantelyat A, Roemmich RT, Chib VS (2023) Dopamine facilitates the translation of physical exertion into assessments of effort. npj Parkinson’s Disease 9:51.

38. Pageaux B (2014) The psychobiological model of endurance performance: An effort-based decision-making theory to explain self-paced endurance performance. Sports Medicine 44:1319–1320.

39. Pageaux B and Lepers R (2016) Fatigue induced by physical and mental exertion increases perception of effort impairs subsequent endurance performance. Frontiers in Psychology 7:1–9.

40. Pierce J, Gray JR, Simpson S, MacAskill M, Höchenberger R, Sogo H, Lindeløv JK (2019) PsychoPy2: Experiments in behavior made easy. Behavioral Research Methods 51:195–203.

41. Pessiglione M, Vinckier F, Bouret S, Daunizeau J, Le Bouc R (2018) Why not try harder? Computation approach to motivation deficits in neuro-psychiatric diseases. Brain 141:629–650.

42. R Core Team (2021) R: A language and environment for statistical computing. R Foundation for Statistical Computing, Vienna, Austria. URL https://www.R-project.org/.

43. Shenhav A, Musslick S, Lieder F, Kool W, Griffiths TL, Cohen JD, Botvinick MM (2017) Toward a rational mechanistic account of mental effort. Annual Review of Neuroscience 40:99–124.

44. Steward G and Chib VS (2024) The neurobiology of cognitive fatigue and its influence on effort-based choice. bioRxiv 2024.07.15.603598. 10.1101/2024.07.15.603598.

45. Umesh A, Kutten KS, Hogan PS, Ratnanather JT, Chib VS (2020) Motor cortical thickness is related to effort-based decision-making in humans. Journal of Neurophysiology 123:2373–2381.

46. Westbrook A, Kester D, Braver TS (2013) What is the subjective cost of cognitive effort? Load, trait, and aging effects revealed by economic preference. PLOS One 8:e68210.

47. Westbrook A and Braver TS (2015) Cognitive effort: A neuroeconomic approach. Cognitive, Affective, and Behavioral Neuroscience 14:395–415.

48. Westbrook A, Lamichhane B, Braver T (2019) The subjective value of cognitive effort is encoded by a domain-general valuation network. Journal of Neuroscience 39:3934–3947.

49. Wylie GR, Yao B, Genova HM, Chen MH, DeLuca J (2020) Using functional connectivity changes associated with cognitive fatigue to delineate a fatigue network. Scientific Reports 10:21927–21938.

