## Supplementary Material for "Neural mechanisms underlying the effects of cognitive fatigue on physical effort-based choice"

**SUPPLEMENTARY FIGURES**

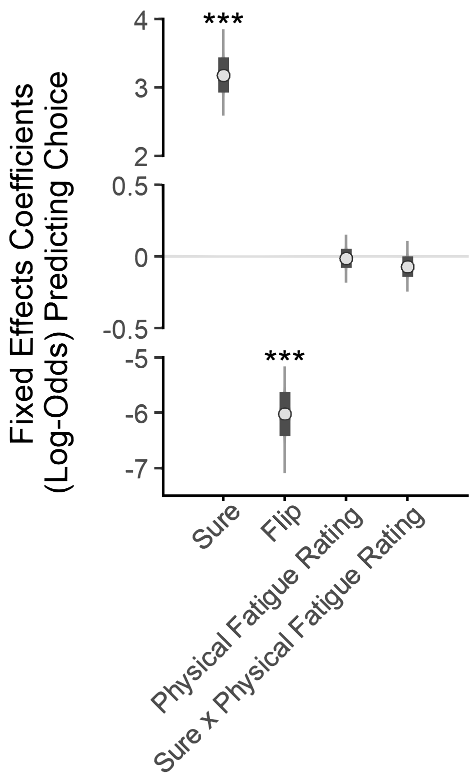


**Supplementary Figure 1 (n = 25)**

Bayesian hierarchical logistic regression predicting choices to select the risky or sure options during the fatigue choice phase. There is no main effect of physical fatigue rating on choice nor a significant interaction between physical fatigue rating and the value of the sure option. The asterisks show significant regressors (*: $p < 0.05$; **: $p < 0.01$; ***: $p < 0.001$). Bars are SEM and lines are 95% confidence intervals. Model results are stable regardless of the broad prior used.

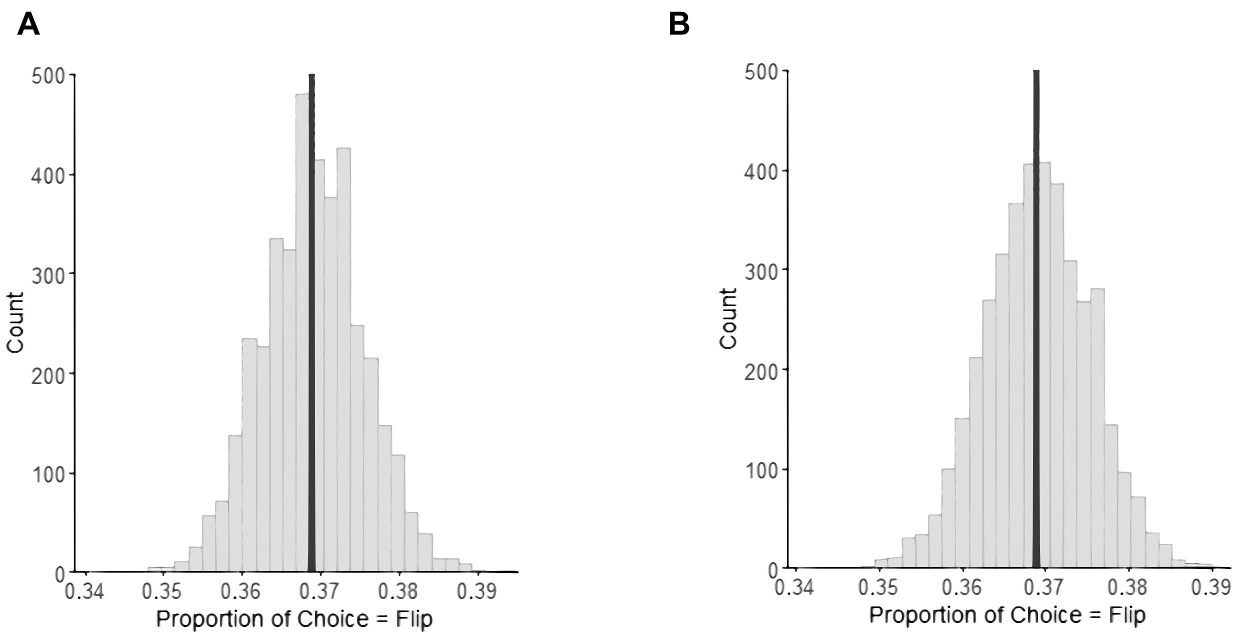


**Supplementary Figure 2 (n = 25)**

**A, B**, Posterior predictive checks of the cognitive fatigue rating (left) and physical fatigue rating (right) behavioral choice models used in our analysis. The centered, vertical black line is the sample proportion of choice scenarios in which participants chose the flip option. The shaded columns represent the proportions predicted by random draws from the posterior distribution of our models. Predicted proportions immediately surrounding the actual proportion in our data have higher counts, demonstrating that our models accurately simulate the choice data.

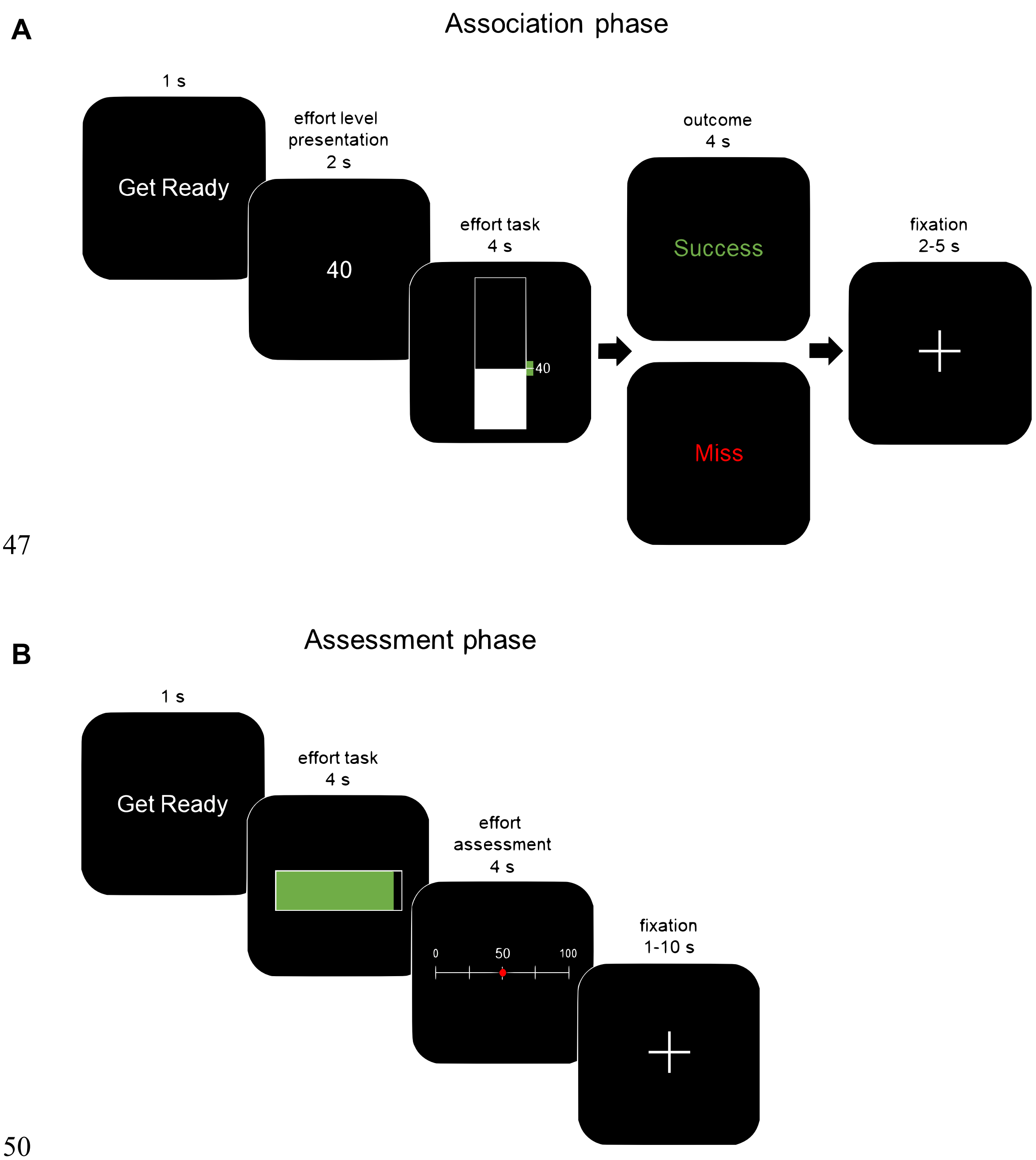


**Supplementary Figure 3**

**A**, Association phase; participants were trained to associate numeric effort levels with force exerted on a hand-clench dynamometer. Effort levels ranged from 0 (no force) to 100 (80% of MVC). A training block consisted of five trials, and there were training blocks for effort levels 10 to 80 in increments of 10 effort levels. Each trial began with the presentation of a numeric target, followed by an effort task with real-time visual feedback of the exerted force represented as a bar that increased in height with increased exertion. A target zone was also displayed, turning red to green if participants exerted the target effort. Feedback of success or failure was provided at the end of each trial.

**B**, Assessment phase; participants were instructed to fill a horizontal bar by exerting force on the hand dynamometer. On each trial, a full bar corresponded to a different target effort level that was not revealed to participants. Successfully producing the target effort resulted in the bar turning red to green. Following exertion, participants used a mouse to select a value along a 0-100 number line to indicate the effort level they believed they had just squeezed. No feedback was provided as to the accuracy of participants' reported effort levels.

### SUPPLEMENTARY TABLES

| Prior | Parameter | β | σ | 95% CI | $\hat{R}$ | Bulk ESS |
|---|---|---|---|---|---|---|
| $N(0,1)$ | Intercept | -1.94 | 0.69 | [-3.18, -0.53] | 1.01 | 637 |
| | $EV_{Sure}$ | 2.42 | 0.39 | [1.53, 3.09] | 1.00 | 1,064 |
| | $EV_{Flip}$ | -4.43 | 0.64 | [-5.47, -2.98] | 1.00 | 764 |
| | Fatigue Rating | -0.09 | 0.11 | [-0.33, 0.13] | 1.00 | 4,135 |
| | $EV_{Sure} * Fatigue\ Rating$ | -0.27 | 0.10 | [-0.48, -0.08] | 1.00 | 5,267 |
| $N(0,10)$ | Intercept | -3.04 | 0.63 | [-4.33, -1.81] | 1.00 | 612 |
| | $EV_{Sure}$ | 3.28 | 0.38 | [2.56, 4.08] | 1.00 | 1,236 |
| | $EV_{Flip}$ | -6.17 | 0.63 | [-7.53, -5.04] | 1.00 | 1,005 |
| | Fatigue Rating | -0.09 | 0.11 | [-0.31, 0.12] | 1.00 | 3,487 |
| | $EV_{Sure} * Fatigue\ Rating$ | -0.31 | 0.10 | [-0.51, -0.11] | 1.00 | 4,558 |
| $N(0,10^6)$ | Intercept | -3.07 | 0.65 | [-4.37, -1.82] | 1.00 | 654 |
| | $EV_{Sure}$ | 3.26 | 0.40 | [2.53, 4.15] | 1.00 | 1,248 |
| | $EV_{Flip}$ | -6.22 | 0.64 | [-7.54, -5.07] | 1.00 | 1,064 |
| | Fatigue Rating | -0.09 | 0.11 | [-0.31, 0.12] | 1.00 | 3,948 |
| | $EV_{Sure} * Fatigue\ Rating$ | -0.30 | 0.11 | [-0.51, -0.10] | 1.00 | 5,438 |
| $U(1,\infty)$ | Intercept | -3.05 | 0.66 | [-4.34, -1.77] | 1.01 | 836 |
| | $EV_{Sure}$ | 3.27 | 0.39 | [2.55, 4.05] | 1.00 | 1,484 |
| | $EV_{Flip}$ | -6.21 | 0.62 | [-7.48, -5.07] | 1.00 | 1,328 |
| | Fatigue Rating | -0.09 | 0.11 | [-0.32, 0.13] | 1.00 | 3,219 |
| | $EV_{Sure} * Fatigue\ Rating$ | -0.31 | 0.10 | [-0.51, -0.11] | 1.00 | 4,596 |

**Supplementary Table 1 (n = 25).** Estimates of the posterior distributions for fixed effects parameters in each model that included cognitive fatigue rating as a regressor. The model used for our analysis is outlined in bold. βs and σs are the mean and standard deviation of a given parameter's posterior distribution. The 95% credible intervals, convergence diagnostics ($\hat{R}$), and sampling efficiency (Bulk ESS) are also shown. Model results are stable regardless of the broad prior used.

| Prior | Parameter | β | σ | 95% CI | $\hat{R}$ | Bulk ESS |
|---|---|---|---|---|---|---|
| $N(0,1)$ | Intercept | -1.91 | 0.67 | [-3.09, -0.40] | 1.00 | 597 |
| | $EV_{Sure}$ | 2.37 | 0.41 | [1.37, 3.06] | 1.00 | 912 |
| | $EV_{Flip}$ | -4.38 | 0.65 | [-5.42, -2.80] | 1.00 | 707 |
| | Fatigue Rating | 0.00 | 0.10 | [-0.19, 0.19] | 1.00 | 4,723 |
| | $EV_{Sure} * Fatigue\ Rating$ | -0.06 | 0.11 | [-0.27, 0.15] | 1.00 | 5,055 |
| $N(0,10)$ | Intercept | -2.94 | 0.63 | [-4.21, -1.73] | 1.00 | 961 |
| | $EV_{Sure}$ | 3.19 | 0.39 | [2.48, 3.99] | 1.00 | 1,675 |
| | $EV_{Flip}$ | -6.05 | 0.59 | [-7.30, -4.97] | 1.00 | 1,219 |
| | Fatigue Rating | -0.01 | 0.10 | [-0.21, 0.18] | 1.00 | 3,555 |
| | $EV_{Sure} * Fatigue\ Rating$ | -0.07 | 0.11 | [-0.28, 0.14] | 1.00 | 4,316 |
| $N(0,10^6)$ | Intercept | -3.00 | 0.64 | [-4.30, -1.74] | 1.01 | 679 |
| | $EV_{Sure}$ | 3.21 | 0.38 | [2.51, 3.99] | 1.00 | 1,058 |
| | $EV_{Flip}$ | -6.10 | 0.61 | [-7.40, -5.00] | 1.00 | 910 |
| | Fatigue Rating | -0.01 | 0.10 | [-0.21, 0.18] | 1.00 | 3,800 |
| | $EV_{Sure} * Fatigue\ Rating$ | -0.07 | 0.11 | [-0.29, 0.14] | 1.00 | 4,451 |
| $U(1,\infty)$ | Intercept | -3.00 | 0.61 | [-4.25, -1.81] | 1.02 | 629 |
| | $EV_{Sure}$ | 3.20 | 0.38 | [2.53, 3.99] | 1.01 | 1,105 |
| | $EV_{Flip}$ | -6.10 | 0.60 | [-7.36, -5.00] | 1.01 | 1,101 |
| | Fatigue Rating | -0.01 | 0.10 | [-0.22, 0.18] | 1.00 | 4,143 |
| | $EV_{Sure} * Fatigue\ Rating$ | -0.06 | 0.11 | [-0.27, 0.16] | 1.00 | 3,875 |

**Supplementary Table 2 (n = 25).** Estimates of the posterior distributions for fixed effects parameters in each model that included physical fatigue rating as a regressor. The model used for our analysis is outlined in bold. βs and σs are the mean and standard deviation of a given parameter's posterior distribution. The 95% credible intervals, convergence diagnostics ($\hat{R}$), and sampling efficiency (Bulk ESS) are also shown. Model results are stable regardless of the broad prior used.

| S | F | S | F | S | F |
|---|---|---|---|---|---|
| 5.0 | 10.0 | 16.25 | 40.0 | 27.5 | 70.0 |
| 5.0 | 17.5 | 16.25 | 47.5 | 27.5 | 77.5 |
| 5.0 | 25.0 | 16.25 | 55.0 | 31.25 | 10.0 |
| 5.0 | 32.5 | 16.25 | 62.5 | 31.25 | 17.5 |
| 5.0 | 40.0 | 16.25 | 70.0 | 31.25 | 25.0 |
| 5.0 | 47.5 | 16.25 | 77.5 | 31.25 | 32.5 |
| 5.0 | 55.0 | 20.0 | 10.0 | 31.25 | 40.0 |
| 5.0 | 62.5 | 20.0 | 17.5 | 31.25 | 47.5 |
| 5.0 | 70.0 | 20.0 | 25.0 | 31.25 | 55.0 |
| 5.0 | 77.5 | 20.0 | 32.5 | 31.25 | 62.5 |
| 8.75 | 10.0 | 20.0 | 40.0 | 31.25 | 70.0 |
| 8.75 | 17.5 | 20.0 | 47.5 | 31.25 | 77.5 |
| 8.75 | 25.0 | 20.0 | 55.0 | 35.0 | 10.0 |
| 8.75 | 32.5 | 20.0 | 62.5 | 35.0 | 17.5 |
| 8.75 | 40.0 | 20.0 | 70.0 | 35.0 | 25.0 |
| 8.75 | 47.5 | 20.0 | 77.5 | 35.0 | 32.5 |
| 8.75 | 55.0 | 23.75 | 10.0 | 35.0 | 40.0 |
| 8.75 | 62.5 | 23.75 | 17.5 | 35.0 | 47.5 |
| 8.75 | 70.0 | 23.75 | 25.0 | 35.0 | 55.0 |
| 8.75 | 77.5 | 23.75 | 32.5 | 35.0 | 62.5 |
| 12.5 | 10.0 | 23.75 | 40.0 | 35.0 | 70.0 |
| 12.5 | 17.5 | 23.75 | 47.5 | 35.0 | 77.5 |
| 12.5 | 25.0 | 23.75 | 55.0 | 38.75 | 10.0 |
| 12.5 | 32.5 | 23.75 | 62.5 | 38.75 | 17.5 |
| 12.5 | 40.0 | 23.75 | 70.0 | 38.75 | 25.0 |
| 12.5 | 47.5 | 23.75 | 77.5 | 38.75 | 32.5 |
| 12.5 | 55.0 | 27.5 | 10.0 | 38.75 | 40.0 |
| 12.5 | 62.5 | 27.5 | 17.5 | 38.75 | 47.5 |
| 12.5 | 70.0 | 27.5 | 25.0 | 38.75 | 55.0 |
| 12.5 | 77.5 | 27.5 | 32.5 | 38.75 | 62.5 |
| 16.25 | 10.0 | 27.5 | 40.0 | 38.75 | 70.0 |
| 16.25 | 17.5 | 27.5 | 47.5 | 38.75 | 77.5 |
| 16.25 | 25.0 | 27.5 | 55.0 | | |
| 16.25 | 32.5 | 27.5 | 62.5 | | |

**Supplementary Table 3.** Effort choices presented during the choice phases. Effort level pairings that comprise the 100 risky choices presented to participants in a random order over the course of the baseline choice phase, and a pseudo-random order over the course of the fatigue choice phase. Each choice consists of a sure (S) option promising lower exertion with certainty, and a risk (F) option with higher potential exertion with uncertainty.
